# Tracking the dynamics of perisaccadic visual signals with magnetoencephalography

**DOI:** 10.1101/2022.01.19.476936

**Authors:** Konstantinos Nasiotis, Sujaya Neupane, Shahab Bakhtiari, Sylvain Baillet, Christopher C. Pack

## Abstract

Many brain functions are difficult to localize, as they involve distributed networks that reconfigure themselves on short timescales. One example is the integration of oculomotor and visual signals that occurs with each eye movement: The brain must combine motor signals about the eye displacement with retinal signals, to infer the structure of the surrounding environment. Our understanding of this process comes primarily from single-neuron recordings, which are limited in spatial extent, or fMRI measurements, which have poor temporal resolution. We have therefore studied visual processing during eye movements, using magnetoencephalography (MEG), which affords high spatiotemporal resolution. Human subjects performed a task in which they reported the orientation of a visual stimulus while executing a saccade. After removal of eye movement artifacts, time-frequency analysis revealed a signal that propagated in the beta-frequency band from parietal cortex to visual cortex. This signal had the characteristics of perisaccadic “remapping”, a neural signature of the integration of oculomotor and visual signals. These results reveal a novel mechanism of visual perception and demonstrate that MEG can provide a useful window into distributed brain functions.

## I. Introduction

Many brain functions are localized to specific cortical regions. As a result, punctate lesions can create highly specific sensory or motor deficits, such as an inability to perceive faces (prosopagnosia) or an inability to speak (aphasia). But many other brain functions rely on distributed processing, even for simple behaviors. A well-known example is the integration of visual and oculomotor signals that supports accurate spatial vision. This kind of integration is necessary because humans move their eyes several times per second, and each eye movement introduces a dramatic disturbance of vision, as the retinal image is abruptly displaced.

The brain compensates for these disturbances by updating retinal signals according to copies of the oculomotor commands (Hemholtz, 1925; Wurtz, 2008). These *corollary discharge* signals arise in the brainstem, after which they are relayed to the frontal cortex and distributed to parietal and occipital regions concerned with vision (Wurtz, 2008). In contrast, the visual signals themselves arise in the retina, are relayed through the thalamus, and ultimately reach the occipital lobe and its projection targets throughout the cortex. Within these distributed cortical networks, visual and oculomotor signals are combined in such a way as to support accurate spatial vision. At present, very little is known about how the brain performs this integration(Neupane et al., 2017).

Most of our knowledge about perisaccadic vision comes from single-neuron studies in non-human primates (Neupane et al., 2020; Wurtz, 2008). In these studies, individual neurons respond to visual stimuli at specific positions, but they alter their encoding of space when an eye movement is being planned (Duhamel et al., 1992; Neupane et al., 2016a,b; Sommer & Wurtz, 2006; Nakamura & Colby, 2002). This *perisaccadic remapping* is thought to play a variety of roles in perception (Cicchini et al., 2013), memory (Umeno & Goldberg, 2001), and learning (Laamerad et al., 2020).

Studies of remapping in single neurons necessarily provide a very limited window into the distributed operations that occur throughout the brain during eye movements. Brain imaging experiments using fMRI have therefore been designed to obtain a more comprehensive spatial view. However, fMRI has a limited temporal resolution (Lescroart et al., 2016; Merriam et al., 2007), so that it is unable to precisely track the effects of remapping, which typically endure for only a fraction of a second (Kusunoki & Goldberg, 2003). One way to overcome these limitations is to use magnetoencephalography (MEG), which affords high spatiotemporal resolution (Nasiotis et al., 2017). MEG is capable of resolving activity changes in different brain regions, with temporal resolution that is sufficient to capture rapid changes that occur during eye movements.

We have therefore studied perisaccadic remapping with MEG. After removing eye movement artifacts from the MEG signals, we identified a neural signal that reflected the visual and oculomotor properties of perisaccadic remapping. This signal was localized to the beta frequency band (20-40Hz) and appeared to originate in the parietal cortices and propagate backward through the visual cortex, before arriving in lateral-occipital cortex. This finding suggests that perisaccadic remapping makes use of cortical feedback pathways that are similar to those typically associated with voluntary attention (Cavanagh et al., 2010; Rolfs et al., 2011).

## II. Experimental Methods

### A. Participants and imaging

Data were recorded from 8 healthy, right-handed participants, all of whom had normal or corrected to normal vision. All participants gave written consent prior to participation in the study, which involved a structural MRI, followed by MEG imaging. The experimental protocols were approved by the Research Ethics Board of the Montreal Neurological Institute.

Each participant first underwent an MRI scan, during which they were positioned on their backs with a 32-channel surface coil centered over the occipital pole. Three-dimensional, T1-weighted anatomical MR image volumes covering the entire brain were acquired on a Siemens TIM Trio scanner (3D-MPRAGE, TR/TE= 2300/2.98 ms, TI = 900 ms, 176 sagittally oriented slices, slice thickness = 1 mm, 256 × 240 acquisition matrix).

MEG data were then recorded using a 275-channel (axial gradiometers), whole-head MEG system (CTF MEG International Services Ltd.). Each participant’s head was digitized (typically 200 points) with a 6 degree-of-freedom digitizer (Patriot - Polhemus) prior to MEG data collection. This was used to mark the scalp, eyebrows and nose, and to optimize co-registration with the anatomical MRI. Three head positioning coils were attached to fiducial anatomical locations (nasion, left/right pre-auricular points) to track head movement inside the MEG. Eye movements and blinks were recorded using 2 bipolar electro-oculographic (EOG) channels. EOG leads were placed above and below one eye (vertical channel) and the second channel was placed laterally to the two eyes (horizontal channel). Heart activity was recorded with one channel (ECG), with electrical reference at the opposite clavicle, for subsequent MEG artifact detection and removal. All data were sampled at 2400 Hz.

During the MEG imaging, visual stimuli were presented on a screen placed in front of the participants at a viewing distance of 45 cm, which permitted visual stimulation up to 25×20 degrees of eccentricity. The display system consisted of a projector (VPixx Technologies, PROPPixxx) located outside the magnetically shielded room and three reflecting mirrors that directed images to the screen. The refresh rate of the projector was 120 Hz with a resolution of 1920×1080 pixels.

### B. Experimental task

Participants were seated in a dimly illuminated room (0.13 cd/m^2^) and asked to fixate on one of two possible red dots of 0.3 degrees radius. After a random (500 – 1500 ms) delay, a white square probe (P1; 4 deg. across, 34.6 cd/m^2^) appeared for 50 ms at a random position in the visual field, allowing us to map the retinotopic organization of each MEG voxel (Nasiotis et al., 2017). The target was subsequently displaced by 10 degrees horizontally (Figure 1), and participants were instructed to perform a saccade to reacquire fixation.

**Figure 1.**
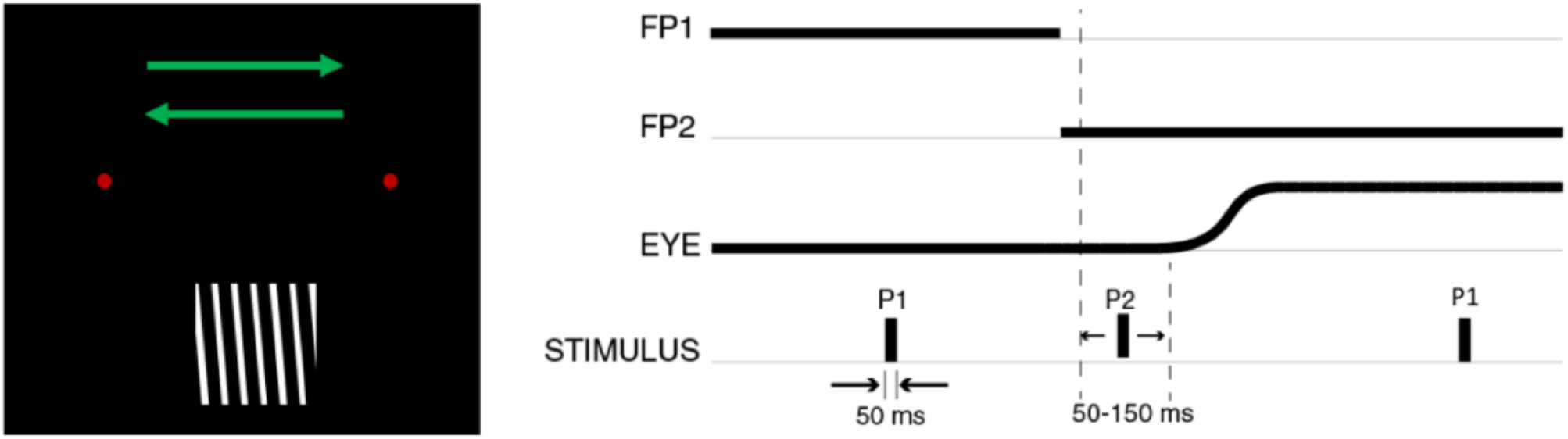
Experimental design. Subjects were asked to perform saccades between two targets (red dots) while a probe appeared elsewhere in the visual field. The timing of the probe could be during fixation (P1) or close to saccade onset (P2). P1 probes were high-contrast white squares (not shown), while P2 probes contained orientation lines (tilted 5 degrees left/right). On each trial, subjects were asked to report the orientation of the P2 probes through a button press (depicted on the left of the figure).

Around the onset of the saccade, a second probe (P2) was presented for 50 ms in the lower visual field, midway between the two targets. The timing of this probe was adjusted according to each participant’s typical saccade latency, so as to occur around the onset of each saccade; typical timing was 50 – 150 ms after the onset of the second fixation target. This probe was in the form of an oriented, high-contrast grating, tilted by 5° clockwise or counter clockwise from vertical. After completion of the saccade, participants were asked to report the orientation of the P2 probe relative to vertical (left or right), via a button press. If no response was given by the participant within five seconds, a new trial was initiated. Feedback was given after each response by turning the fixation dot green or gray, for correct and failed trials respectively.

Monitoring of the probes’ on and off states was performed by a photodiode that was located at the corner of the screen, hidden from the participant’s visual field. A P2 trial was considered successful only when the saccade was initiated after the probe was off, and not later than 200ms after the probe offset. The photodiode was sampled from the acquisition system at the same rate as the MEG signals.

At the end of the experiment, every participant also participated in a 10-minute experimental run, in which the same saccades were executed in the absence of any visual probes. These trials were used during the analysis for baseline correction.

### C. Detection and removal of eye movement artifacts

As with electroencephalography (EEG), MEG signals are susceptible to artifacts produced by eye movements. To detect these artifacts, we used independent component analysis (ICA) with the InfoMax algorithm (Bell & Sejnowski, 1995), in conjunction with the natural gradient feature of (Amari, 2009)that is integrated within EEGLAB (Delorme & Makeig, 2004).

Specifically, we consider the matrix of sensor outputs **X**, where each raw vector represents a different sensor **X**_i_ = [x_1_, x_2,_ … x_k_], with i ∈ [1, m] for m sensors. This matrix can be characterized as the product of an m x m mixing matrix **A** and a series of sources **S**, such as:

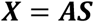

ICA tries to find the unmixing matrix W, that isolates the sources that contributed to the observed matrix **X**:

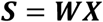

This is achieved by minimizing the mutual information between contributing sources (or in other words, searching for components that maximize their independence). Mutual information is given by:

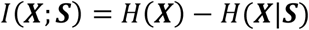

Here *H*(***X***|***S***) =*H*(***X, S***) − *H*(***S***) is the conditional entropy and *H*(***X***) is the entropy of **X**. The entropy is given by:

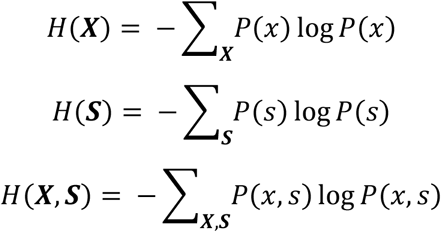

Where P(x) is the probability of observing x in **X** and P(x,s) is the joint probability of x and s.

The InfoMax algorithm that is used for computing **W** consists of the following steps (Langlois, 2010):

1. Initialize ***W***_0_ with random values.
2. 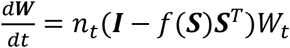
3. If |*max*(*d****W***_***i***,***j***_)| > *ε for i,j* **∈** m, repeat from step 2

Here f(**Y**) = tanh(**Y**), ***I*** is an *m x m* identity matrix, *n*_*t*_ is a learning rate variable, and *ε* is a convergence threshold.

Rejection of components was performed by zeroing out the rows in **S**.

### D. Time-Frequency analysis

Our main goal was to identify MEG signals that correlated with perisaccadic remapping, and to localize them in frequency and time. To this end, we used the complex Morlet wavelet, which has point spread functions with Gaussian shapes in both time (temporal resolution) and in frequency (spectral resolution). Resolution is given in units of the FWHM (full width half maximum) of the Gaussian.

Dilations and translations of the “Mother function,” or “analyzing wavelet” Φ(t), define an orthogonal basis, or wavelet basis:

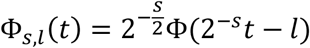

The mother wavelet *Φ(t)* defines an orthogonal basis through dilations and translations (s and l respectively). Therefore, a signal can be decomposed into wavelets through:

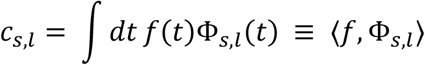

where *f(t)* is the signal that is decomposed into wavelets, and the *c*_*s,l*_ are the wavelet coefficients.

All trials were segmented around the timing of the saccade offset, for both saccades to the left and to the right (−1000:1000 ms), and wavelet decompositions were performed for frequencies between [6-90] Hz. The mother wavelet was selected with central frequency of 1 Hz and time resolution (FWHM) of 3 s. Wavelets that corresponded to each condition were averaged for each participant.

### E. ERSD analysis

To detect modulations in MEG signal power, we used event related synchronization / desynchronization (ERSD) analysis. ERSD quantifies the spectral modulation of a signal during a post event period relative to a baseline period.

ERSD is given by the formula:

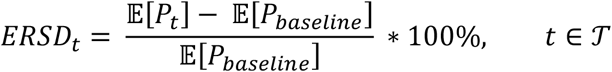

where P_t_ denotes the signal power within a frequency band during the event related period 𝒯, and P_baseline_ is the averaged power within the baseline selected:

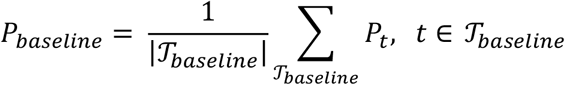

By convention, positive values are referred to as event related synchronization (ERS) and negative values as event related desynchronization (ERD).

ERSD metrics were computed for each wavelet average, with respect to a baseline, which was defined as the period [−800, −200] ms before the saccade offset for all frequency bins. All participants’ cortical responses were projected, rectified, and smoothed (3mm smoothing) on the MNI/ICBM152 average brain template (Fonov et al., 2009). Each participant’s ERSDs were averaged across all runs. To quantify the perisaccadic power modulation in the ROI corresponding to the parietal cortex, ERSDs of all subjects were averaged.

To detect the spatial statistical significance of ERSD events, signals from each cortical source were band-pass filtered and subjected to a paired permutation test (saccades with a probe/without a probe - 1000 randomizations). The statistical significance threshold was set to α = 0.05 (uncorrected), and a separate test was performed for every time sample.

## III. RESULTS

Single-neuron recordings have revealed an eye movement input that triggers remapping of visual receptive fields (Duhamel et al., 1992; Heiser et al., 2005; Neupane et al., 2016a; Sommer & Wurtz, 2006; Umeno & Goldberg, 1997). However, the circuitry that supports the integration of oculomotor and visual signals is not well understood. Previous work suggests a role for oscillatory brain activity in this function (Bosman et al., 2012; Neupane et al., 2017; Zanos et al., 2015), so we used MEG to track the flow of oculomotor influences across visual cortex, in frequency and in time.

### A. Effectiveness of eye movement artifact removal from MEG signals

One barrier to the use of MEG or EEG in studies of eye movements is that the eye musculature acts as dipole, which generates electromagnetic sinks and sources independent of and significantly larger than signals arising from brain activity. As described in the Methods, we therefore implemented an ICA-based method for rejecting eye movement artifacts. We first demonstrate the effectiveness of this approach and then describe the main scientific results.

As described below, human participants performed a task that required fixation on a red dot that appeared in one of two fixed positions on the lateral axis. Displacement of the red dot provided an instruction to make a horizontal eye movement. Eye movements were monitored via two electrooculogram (EOG) channels that detected both horizontal and vertical eye movements, providing the ICA components with a ground truth signal to constrain artifact removal. MEG signals were measured from a 273 channel CTF system.

ICA was performed on the continuous raw signals, and a single component that tracked the lateral movement of the eyes was detected and successfully removed from the recordings. Cross correlation between the eye signal recorded with EOG and each ICA component showed a single component with high correlation (p<<.001) at zero lag. Visual inspection confirmed that the components with high correlation matched either saccadic eye movement or blinks. The spatial topography of this component showed that it originated from the frontal sensors, as expected (Figure 2, right). Thus, ICA was able to precisely filter out eye movement artifacts. Furthermore, although blinks originate from the same physiological source, ICA was able to differentiate blinks and lateral eye movements into different components (Figure 2).

**Figure 2.**
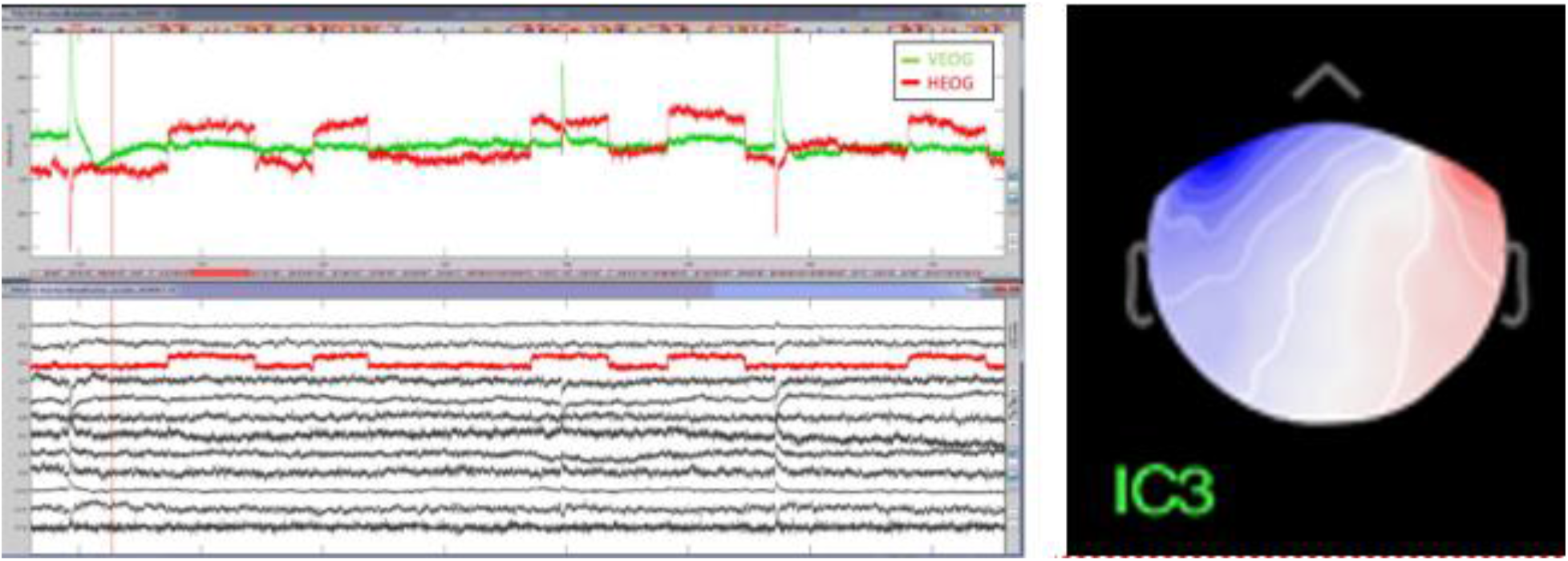
ICA Analysis components. Left: Top row: Electrooculography signals (EOG). The large deflections on the vertical EOG (VEOG) channel reflect eye blinks. Since the saccades were performed on the horizontal axis, the horizontal EOG channel (HEOG) follows the trajectory of the eyes. Bottom row: 20 first MEG ICA components. For all subjects, ICA decomposition revealed a single component that was highly correlated with the HEOG. On this example, component 3 is highlighted to indicate the resemblance to the eye movement. Right: Spatial distribution of coefficients for component 3.

### B. Time-frequency decomposition shows remapping signals in parietal cortex

Having removed eye movement artifacts from the MEG signals, we then characterized oculomotor influences on visual cortical responses. Eight human participants performed a simple task (Figure 1) requiring a horizontal saccade in parallel with an orientation discrimination task (see Methods). The oriented target stimulus was designed in such a way that it appeared in one visual hemifield before each saccade and in the opposite hemifield after the saccade. Consequently, any remapped response could be readily identified, as it would appear in the cortex ipsilateral to the visual stimulus. The paradigm also ensured that participants focused their attention on the target stimulus, which is useful insofar as attention appears to be important for remapping (Cavanagh et al., 2010; Rolfs et al., 2011).

To detect remapped responses to visual stimulus flashed just prior to eye movement, we first focused on a specific ROI defined by parietal cortex. This choice was motivated by the original non-human primate studies showing remapped responses in the lateral intraparietal area (Duhamel). As described in the Methods, we focused on the ERSD response; previous work has shown that ERSD increases are linked to decreased neural activity, and ERSD decreases are linked to increased neural activity (Pfurtscheller & Lopes da Silva, 1999).

Figure 3 shows the ERSD response in the left parietal cortex for a saccade to the left, averaged across participants. As indicated above, this condition should elicit a remapped response to a visual stimulus presented just before saccade onset, as it is remapped to the opposite hemisphere by the impending saccade. Consistent with this idea, a time-frequency analysis (Figure 3, top left) of the parietal MEG sources showed a strong decrease in the ERSD response (blue colors) that was most identifiable in beta band, immediately after the saccade. A similar response appeared in the absence of a visual stimulus (Figure 3, top right), but it was weaker in amplitude.

**Figure 3.**
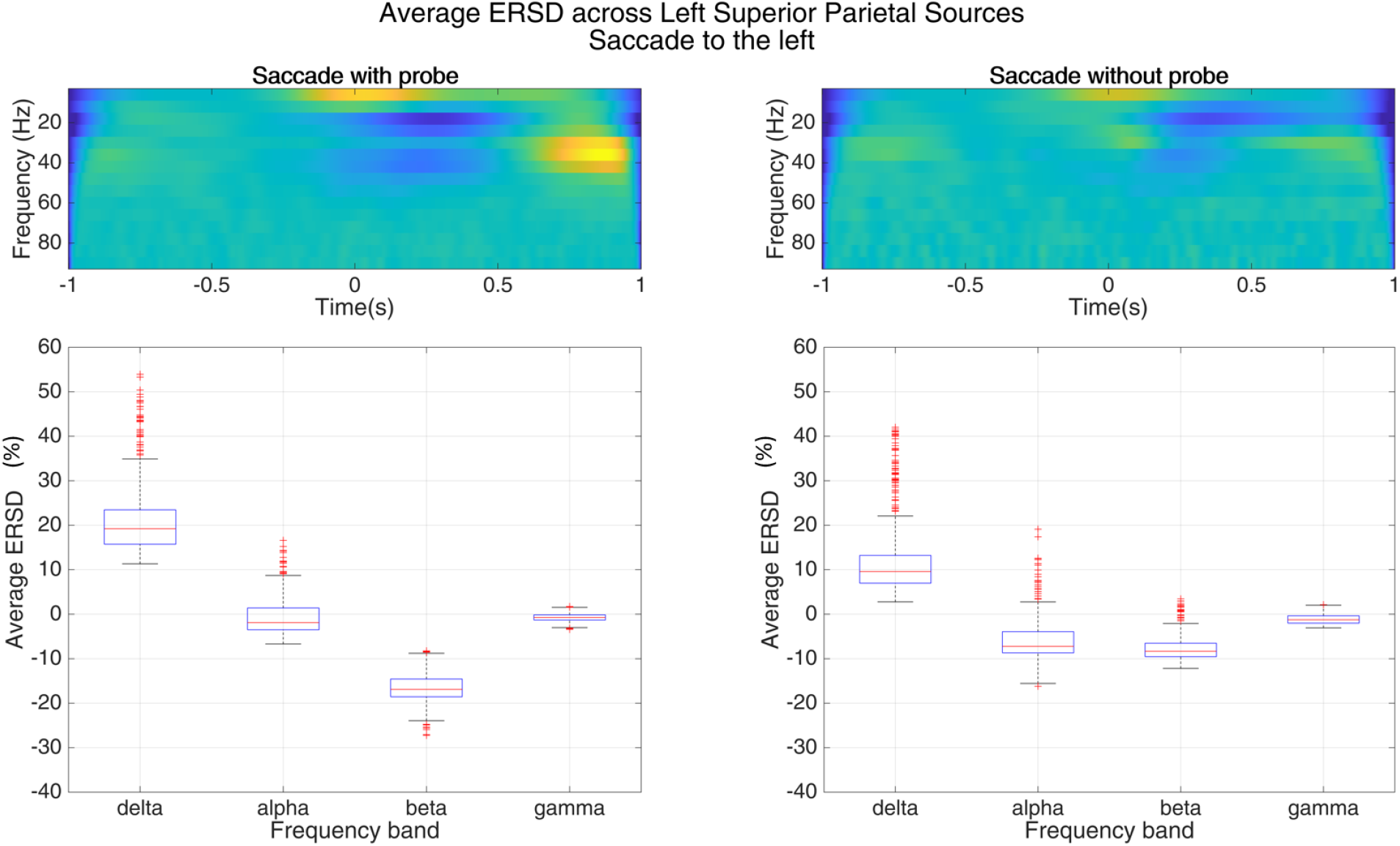
Top row: Averaged perisaccadic ERSD values for left superior parietal sources for a saccade to the left for the conditions where a probe was presented or not. Bottom row: Averaged ERSD values for each frequency band between [0, 500] ms. The beta band shows a significant decrease on the sources that are expected to show remapping when a probe is presented.

To quantify these effects, we examined ERSD response within four standard frequency bands, comparing the probe and no-probe conditions (Figure 3, bottom). Because remapping depends on the presence of a visual stimulus, this contrast served to identify candidate signals. As indicated by the top row of Figure 3, the largest difference between the two conditions occurred for signals in the β-band. We therefore focused on this frequency for subsequent analyses.

To examine these results across the rest of the cortex, we performed a permutation test on each time-sample, which yielded a time-resolved statistical map of the entire cortical surface. Figure 5 shows the resulting map, for both saccade directions, at different time points ranging from −100 ms before each saccade offset to 280 ms after saccade offset. Saccades in the presence of a target probe that appeared just before the saccade caused a decrease of β-band power in the ipsilateral cortex, compared to saccades in the dark, which generally caused an increase at 100 ms after the saccade offset (Figure 4). β-power also decreased significantly after saccade offset, compared to other frequency bands. These results therefore demonstrate the appearance of remapping signals in MEG.

**Figure 4.**
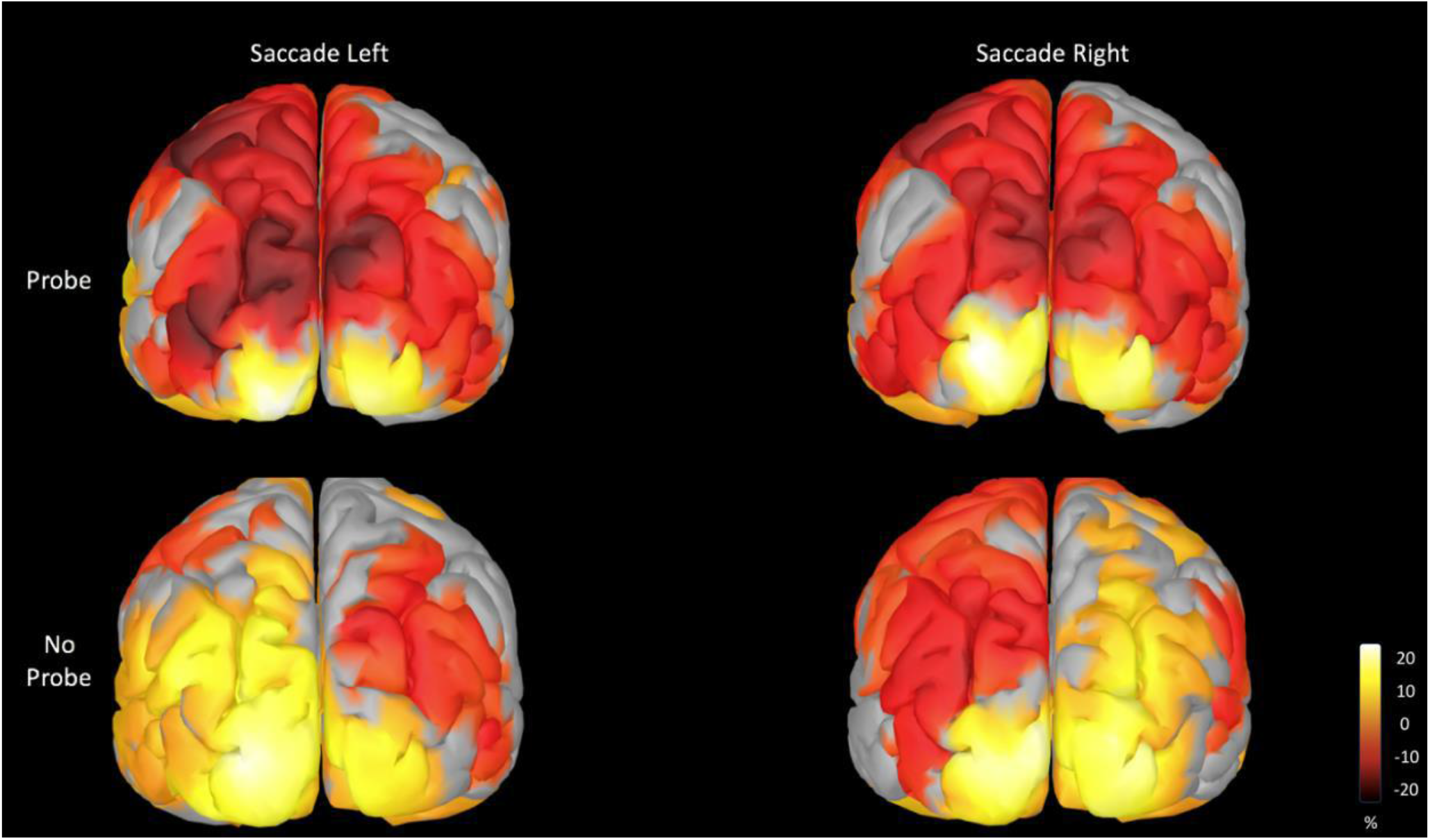
ERSD maps of the probe/no probe conditions for both saccade directions. All figures are synced at 100ms after the saccade offset for the β-band. Power modulation was normalized to a baseline [−800, −200]ms relative to the saccade offset. The presence or the absence of a probe right before the saccade, distinctly affects the β-band activity on the ipsilateral parietal cortex for both saccadic directions.

**Figure 5.**
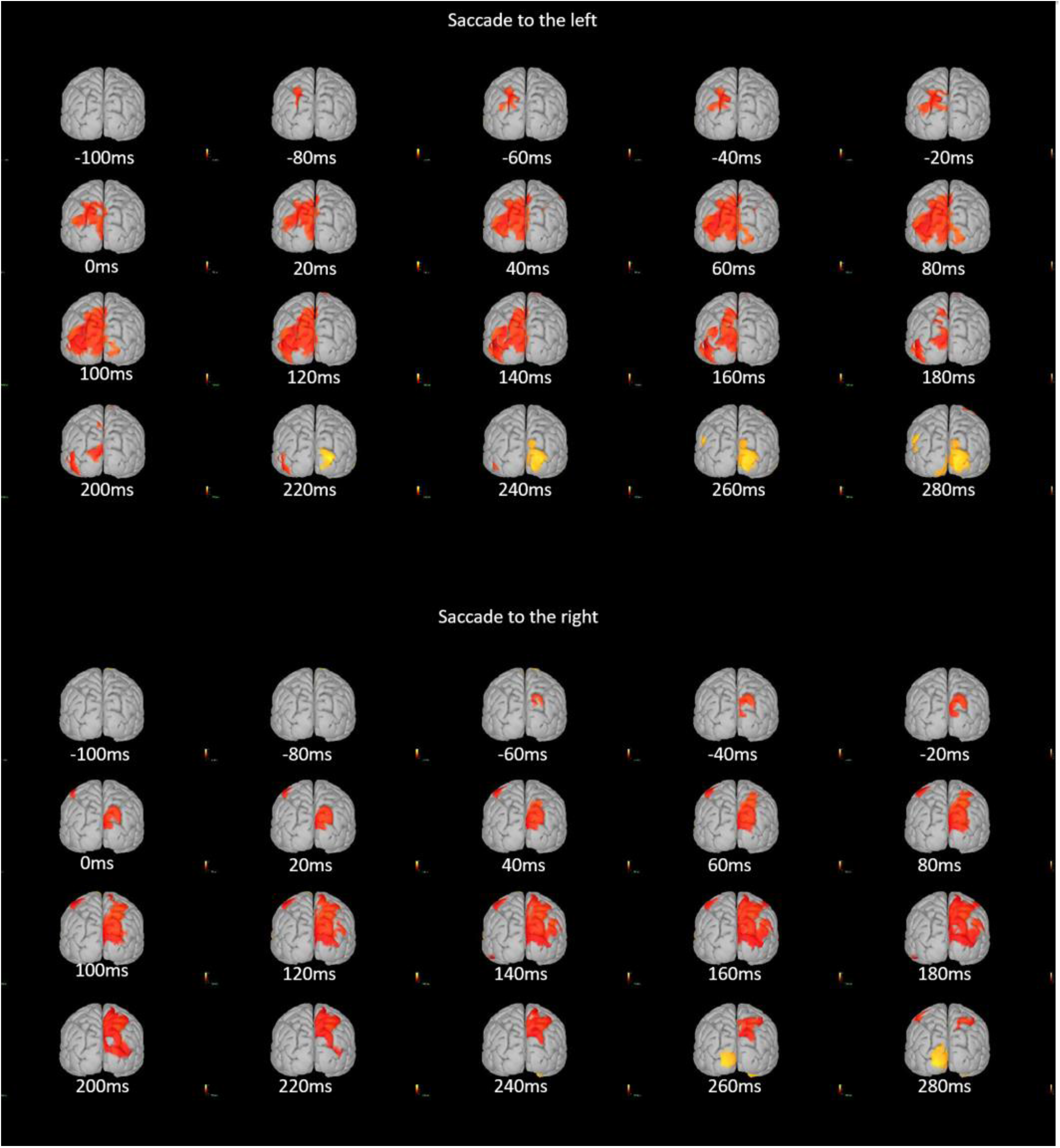
Two-tailed permutation test (1000 repetitions) for all subjects between probe/no probe conditions, for saccades to the left (Top) and to the right (Bottom). The test was performed separately for every time-sample. The figure displays multiple snapshots on different timestamps ranging [-100,280]ms relative to the saccade offset. Due to the selection of the position of the probe, saccades to the left would showcase forward remapping on the left hemisphere (equally saccades to the right, on the right hemisphere). The maps indicate an early decrease of β-power in the ipsilateral superior parietal cortex, followed by an expansion of the decrease towards the inferior parietal and the lateral-occipital cortex. Finally, a sluggish β-band increase in the contralateral lateral-occipital cortex is observed.

### C. Remapped signal propagates across visual hierarchy

The results in the previous section identify a widespread cortical signal related to perisaccadic remapping, localized to the β frequency band and the immediate post-saccadic time period. To infer the possible flow of information within the brain, we next performed a more fine-grained analysis of the temporal progression of remapping signals across cortical areas. As in the previous analysis, we relied on paired permutation tests across time samples for every cortical source, to detect activity that differed significantly between the probe and no-probe conditions.

For this analysis, we focused on ROIs covering the superior parietal, inferior parietal and lateral occipital cortices, in both hemispheres. These ROIs showed significant activation in the analysis of Figure 5 and known to be involved in perisaccadic remapping(Nakamura & Colby, 2002; Wang et al., 2016). Figure 6 shows the sequence of significant remapping responses that appeared in different cortical regions around the time of *leftward* saccades. Recall that remapping responses would be expected to occur in the left hemisphere in this condition.

**Figure 6.**
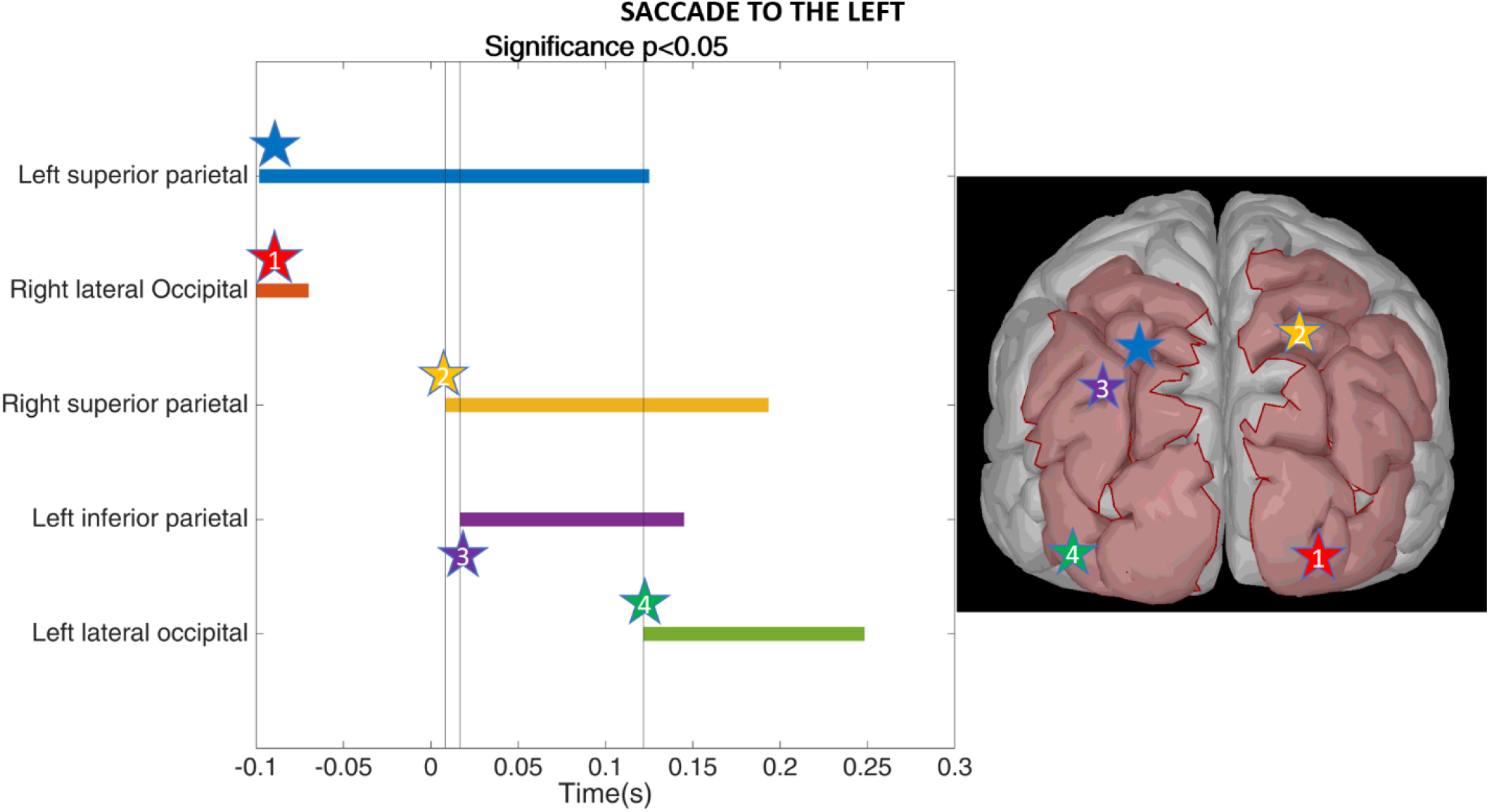
Perisaccadic statistical significance of the centers of clusters within regions of interest for a saccade to the left. The ipsilateral superior parietal cortex shows significance consistently around the saccade.

The results reveal significant β-band suppression in the contralateral visual cortex (Figure 3, red), as expected for an afferent visual response. Similar responses then appear in the contralateral superior parietal cortex (yellow), after which they propagate to the ipsilateral superior parietal cortex (purple) and the ipsilateral visual cortex (green). These ipsilateral responses are consistent with a remapping signal that has previously been detected at the single-neuron level in both parietal (Duhamel et al., 1992; Heiser et al., 2005) and visual (Inaba & Kawano, 2014; Nakamura & Colby, 2002; Neupane et al., 2016a; Yao et al., 2016) cortices. Activity in the left superior parietal lobe (blue) appears before, during, and after the saccade, suggesting a possible role in orchestrating the remapping process.

Together, these results suggest a possible circuit for integrating oculomotor and visual signals. The raw visual information, first detected by the contralateral visual cortex, is subsequently sent to the parietal cortex, where it is combined with an eye movement input. This integrated signal is then relayed across hemispheres (Berman et al., 2005; Heiser et al., 2005; Heiser & Colby, 2006) and fed back to the ipsilateral visual cortex (Neupane et al., 2016a). The latter feedback projection is consistent with previous single-neuron studies supporting a top-down process in remapping (Nakamura & Colby, 2002).

We note also that Figure 5 reveals a very late response to the visual stimulus in the contralateral visual cortex (bottom right, yellow), characterized by an increase in beta-band power. A similar response has been detected in single-neuron recordings (Neupane et al., 2016a, 2020; Tolias et al., 2001).

## IV. Conclusion

We have shown that brain signals related to the phenomenon of perisaccadic remapping can be recovered with MEG, permitting a more comprehensive view of their spatial and temporal distribution throughout the cortex. This approach circumvents the limitations of previous approaches which have relied on single-neuron or fMRI measurements, each of which has limited resolution in space or time. In this regard, the contribution of MEG is critical, because the integration of visual and oculomotor signals takes place on brief time scales and is distributed throughout a network of brain regions.

Our experiment was designed to segregate purely visual from remapped visual signals, by forcing the latter to appear in the ipsilateral hemisphere (Figure 1). In the absence of perisaccadic remapping, strong ipsilateral responses are absent in most of visual cortex and relatively rare in most other brain regions (Arcaro & Livingstone, 2021). With this paradigm, we were able to detect remapped visual responses (Figure 4) and to determine that they likely arise from cross-hemispheric connections in the parietal lobe (Figures 5 and 6). From there they appear to propagate more posteriorly to the occipital cortex (Figure 5).

The importance of feedback for visual perception has long been appreciated from a theoretical standpoint (Bosman et al., 2012; Lee & Mumford, 2003). It is generally thought to be involved in selecting and enhancing the contribution of behaviorally-relevant sensory inputs, and as such it is often associated with voluntary attention. Indeed, attention appears to be particularly important for perisaccadic remapping (Cavanagh et al., 2010; Rolfs et al., 2011).

Consistent with this idea, we found that the remapped signals were strongest in the beta frequency band (Figure 3), which is often implicated in saccades (Zanos et al., 2015) and feedback processing more generally (Bastos et al., 2015). In this regard, it is surprising that we did not detect remapping signals in the frontal cortex; it remains to be seen whether this reflects a limitation of the method or a genuine property of perisaccadic visual processing.

## Acknowledgements

This work was supported by a grant to C.C.P. from the CIHR (MOP-115178). S.B. received funding from NIH (R01 EB026299), a Discovery Grant from the Natural Science and Engineering Research Council of Canada (436355-13), the CIHR Canada Research Chair in Neural Dynamics of Brain Systems, the Brain Canada Foundation with support from Health Canada, and the Innovative Ideas program from the Canada First Research Excellence Fund, awarded to McGill University for the Healthy Brains for Healthy Lives initiative. This research was undertaken thanks in part to funding from the Canada First Research Excellence Fund, awarded to McGill University for the Healthy Brains for Healthy Lives initiative.

## Notes

### Competing Interest Statement

The authors have declared no competing interest.

